# Genome sequencing reveals novel pathogenic deep-intronic *PCDH15* variants, amenable to antisense oligonucleotide-based splice correction

**DOI:** 10.64898/2026.08.20.746067

**Authors:** Kim Rodenburg, Leony Fenwick, Ronald Pennings, Lonneke Haer-Wigman, Tamar Ben-Yosef, Femke van Erp, Janine Reurink, Christian Gilissen, L. Ingeborgh van den Born, Frans P.M. Cremers, Yuval Cohen, Helger Yntema, Erik de Vrieze, Hannie Kremer, Suzanne E. de Bruijn, Rob W.J. Collin, Susanne Roosing, Erwin van Wijk

## Abstract

Despite substantial advances in diagnostic testing, 10-15% of Usher syndrome patients remain without a genetic diagnosis, having significant implications for genetic counseling and potential future therapeutic interventions. In this study, genome sequencing data from probands clinically presenting with Usher syndrome were analyzed. Two novel deep-intronic variants were identified in *PCDH15,* c.3983+3635A>G and c.3123-1728A>G, in two independent patients.

Both deep-intronic variants were classified as likely pathogenic and predicted to alter *PCDH15* pre-mRNA splicing. Using a minigene splice assay and iPSC-derived photoreceptor precursor cells from patients, we confirmed that both variants lead to the inclusion of a pseudoexon in the *PCDH15* transcript introducing a stop codon and subsequent premature termination of protein translation. We designed and evaluated antisense oligonucleotides (ASOs) with the purpose of redirecting aberrant pre-mRNA splicing caused by both deep-intronic variants. For both variants, designed ASOs were successful in restoring normal splicing patterns, highlighting their potential as a future therapeutic intervention strategy to halt the progression of retinitis pigmentosa caused by these novel variants.

Overall, these findings contribute to the understanding of Usher syndrome caused by deep-intronic pathogenic variants in *PCDH15* and describe for the first time the use of an ASO-mediated splice correction strategy for individuals diagnosed with these variants.

## Introduction

Usher syndrome (USH) is a rare autosomal recessive disorder that is typically characterized by bilateral congenital sensorineural hearing loss, bilateral vision loss due to retinitis pigmentosa (RP) and, depending on the clinical type, vestibular dysfunction [1–3]. Based on a prevalence of approximately 1:10,000, an estimated 400,000 individuals suffer from USH [2]. Based on the characteristics of disease, four clinical types are described, with USH type 1 (USH1) being most severe.

Pathogenic variants in so far 11 different genes have been identified to cause Usher syndrome. One of the genes involved in USH1 is *PCDH15* (MIM #605514), which encodes protocadherin 15, a calcium-dependent adhesion protein which belongs to the cadherin superfamily and is essential for the development, maintenance and function of the sensory cells in the inner ear and retina [4, 5]. Pathogenic variants in this gene account for approximately 8-20% of all USH1 cases and are also described to be associated with autosomal recessive non-syndromic hearing loss 23 (DFNB23) [4, 6, 7]. To date, 1456 unique (likely) pathogenic variants have been described for *PCDH15* in the Leiden Open Variant Database (LOVD, accessed August 2026). The clinical outcome depends on the severity and combination of variants. Biallelic missense variants generally result in DFNB23, while combinations with at least one severe variant, such as nonsense variants, deletions and insertions, frameshift variants or variants affecting canonical splice sites, are mostly associated with USH1 [5, 7–9]. Despite extensive efforts in genetic testing using traditional sequencing approaches such as targeted panel sequencing and exome sequencing, 10-15% of all USH cases remain without a conclusive genetic diagnosis [9]. In our local cohort, a part of these individuals carries a mono-allelic pathogenic variant in one of the USH-associated genes, suggesting that a second pathogenic variant may reside in regions which are not typically covered by the methods that are routinely applied.

The broader utilization of whole genome sequencing (WGS) has expanded the capacity of variant discovery and prioritization, revealing previously missed variants in regions not covered in exome sequencing, such as deep-intronic variants that lead to aberrant pre-mRNA splicing, a well-known phenomenon reported for other USH types [10–15].

In this study, we applied WGS analysis to two genetically unresolved USH probands and identified two novel pathogenic deep-intronic variants within the *PCDH15* gene, both of which are predicted to alter pre-mRNA splicing. By performing *in vitro* minigene splice assays for both variants, and by assessing the effect of one variant in patient-derived photoreceptor precursor cells (PPCs), we confirmed the inclusion of two pseudoexons that are both predicted to result in the premature termination of protocadherin 15 translation.

To rescue the deleterious effect of these variants, we developed an antisense oligonucleotide (ASO)-based strategy. Experimental validation demonstrated the efficacy of the ASOs in correcting aberrant pre-mRNA splicing, demonstrating its potential as a future therapeutic strategy for individuals carrying these *PCDH15* variants.

## Material and Methods

The study adhered to the tenets of the Declaration of Helsinki and was approved by the local ethics committees of the Radboud University Medical Center (Nijmegen, The Netherlands), the Rotterdam Eye Hospital (Rotterdam, The Netherlands) and the Hillel Yaffe Medical Center (Hadera, Israel). Written informed consent was obtained from all participating individuals or their guardians prior to DNA analysis and inclusion in this study.

## Genome sequencing

Genomic DNA from two probands was isolated from peripheral blood lymphocytes according to standard procedures. WGS was performed at BGI on a BGISeq500 using 2x 150 base pair (bp) paired-end reads with a minimal median coverage of 30-fold per genome. Sequencing reads were mapped to the Human Reference Genome build GRCh38/hg38 using Burrows-Wheeler Aligner V.0.7814 [16]. Single nucleotide variants (SNVs) and small indels (<50 bp) were called using Genome Analysis Toolkit HaplotypeCaller (Broad Institute); structural variants (SVs) were called using Manta Structural variant Caller, based on read-pair evidence and read-depth evidence; copy number Variants (CNVs) were called using Canvas Copy Number Variant caller, based on read-depth evidence; mobile element insertions were identified using the Mobile Element Locator Tool [17–20]. SNVs, small indels, SVs and CNVs were annotated using an in-house pipeline.

## Variant prioritization and analysis

WGS data were analyzed to identify potential pathogenic variants in inherited retinal disease (IRD)– and USH-associated genes. Coding and non-coding SNVs were selected based on a minor allele frequency <0.01 in the gnomAD population database (all populations, V3.1.2, with additional manual checks in v4.1.1 for selected variants). Rare variants were then prioritized based on protein effect and nonsense, start or stop codon disrupting, frameshift, in-frame deletions and insertions, missense, canonical and intronic splice variants were analyzed in further detail. For missense variants we obtained CADD_PHRED and REVEL scores and only the variants meeting the predefined thresholds of both *in silico* prediction tools (CADD_PHRED: >15, range 0-99; REVEL: >0.3, range 0-1) were selected for further investigation as possibly pathogenic candidates [21, 22]. Variants meeting the threshold of only one tool were retained as lower-priority candidates. For potential splice-altering effects of canonical splice variants, missense, synonymous or intronic variants, SpliceAI delta scores were obtained from SpliceAI (https://spliceailookup.broadinstitute.org/) and variants were considered for further testing using an *in vitro* minigene splice assay if at least two out of the four provided scores (acceptor gain (DS_AG), acceptor loss (DS_AL), donor gain (DS_DG) or donor loss (DS_DL)) provided a delta score of ≥0.2 [23]. Alamut™ Visual plus (v1.7.1) was used to visually inspect the genomic context of the positions predicted by SpliceAI delta scores. Coding SVs were prioritized on a minor allele frequency of <0.01 in the 1000 genome database [24]. Inversions and duplications were only considered potentially pathogenic when disrupting an IRD– or USH-associated gene as proposed by De Bruijn et al. [25].

The newly identified *PCDH15* splice-altering variants were assessed via Sanger sequencing in nine additional probands that were monoallelic for a pathogenic *PCDH15* variant. Segregation analysis was facilitated by the availability of DNA samples from relatives. Primer sequences are listed in **Supplemental Table 1.**

## Minigene splice assays in HEK293T cells

Minigene constructs were generated as previously described [26]. In short, for the selected variants, at least 450 nt of the up– and downstream intronic sequences flanking the predicted pseudoexon resulting from the presence of the variant(s) were amplified from the proband’s genomic DNA and cloned into a pCI-Neo plasmid containing the genomic region of *RHO* encompassing exons 3-5 using Gateway cloning technology (Thermo Fisher Scientific, Carlsbad, CA, USA). Both wildtype fragments and fragments containing the variants were cloned. HEK293T cells were transfected using polyethylenimine (PEI) with either a wildtype or mutant minigene construct and harvested 48 hours post transfection. RNA was isolated using the Nucleospin RNA kit (Machery-Nagel, Düren, Germany) following manufacturer’s instructions. 1 µg of total RNA was used as input for cDNA synthesis using the iScript cDNA synthesis kit (Bio-Rad, Hercules, CA, USA) according to manufacturer’s instructions. RT-PCR analyses were performed using primers for *RHO* exons 3 and 5 to assess the splicing pattern of the *PCDH15* regions of interest. Primers to amplify *ACTB* were used as a loading control. PCR fragments were validated by Sanger sequencing. The assay was performed in two biological replicates. Sequences of all primers used are listed in **Supplemental Table 2**.

## Generation of photoreceptor precursor cells

A fresh heparin blood sample was collected from proband DNA20-15335 (harboring *PCDH15* c.3123-1728A>G) and peripheral blood mononuclear cells were isolated and subsequently reprogrammed into induced pluripotent stem cells through episomal nucleofection as described previously at the Stem Cell Technology Center of the Radboudumc Nijmegen, the Netherlands [27]. Induced pluripotent stem cells were differentiated into PPCs by following the first 30 days of the 2D-to-3D differentiation protocol as described by Gonzalez-Cordero [28].

## Antisense oligonucleotides

ASOs were designed for both *PCDH15* deep-intronic variants following previously published guidelines [29]. In addition, a scrambled oligonucleotide (SON), as well as a 3-nucleotide mismatch ASO (3ntMM ASO) of the best performing ASO per target, were used as negative controls. All ASOs were designed with 2’O-MOE chemical modifications and a fully phosphorothioate backbone.

For transfection in HEK293T cells, ASOs were reconstituted in sterile phosphate buffered saline (PBS) to a stock concentration of 1 mM and diluted to work concentrations of 0.1 mM, 0.05 mM, 0.02 mM and 0.01 mM. For each variant, HEK293T cells were co-transfected at approximately 70% confluency, using 100 µL Opti-MEM™ I (Thermo Fisher Scientific, Carlsbad, CA, USA) and 6 µL Fugene (Promega, Madison, WI, USA), with 1 µL of ASO and 500 ng of either wildtype or mutant minigene construct of the matching variant, in 1 mL culture medium. HEK293T cells were harvested 48 hours post transfection and RNA was isolated using the Nucleospin RNA kit (Machery-Nagel, Düren, Germany) following manufacturer’s instructions.

For photoreceptor precursor cells, ASOs were reconstituted in sterile PBS to a stock concentration of 100 µM and diluted to work solutions of 5 µM, 2.5 µM and 1 µM in culture medium. ASOs and SONs were gymnotically delivered to the PPCs on day 20 of differentiation by replacing culture medium for ASO-containing medium. Cycloheximide (0.1 mg/mL Sigma-Aldrich, St. Louis, MO, USA) was added on day 30 and cells were harvested six hours post addition of cycloheximide in 350 µL RAI buffer (Machery-Nagel, Düren, Germany). RNA was isolated using the Nucleospin RNA kit (Machery-Nagel, Düren, Germany) following the manufacturer’s instructions.

1 µg of total RNA was used as input for cDNA synthesis using either iScript cDNA synthesis kit (Bio-Rad, Hercules, CA, UCA) or SuperScript VILO mastermix (Life Technologies, Carlsbad, CA, USA) according to the manufacturer’s protocol. Diluted cDNA was used as input for PCR (according to standard protocols) to assess the ASO-mediated splice correction. PCR products from transfections in HEK293T cells were sequence-verified by Sanger sequencing. Transfections and delivery of ASOs were performed in duplicate. ASO and primer sequences used are listed in **Supplemental Table 3** and **Supplemental Table 4**.

Semi quantitative fragment analysis was performed using Image Lab (Bio Rad). Lanes were manually assigned to ensure consistent alignment across samples. Fragments were automatically detected. Background correction was performed using lane-based background subtraction.

## Results

### Genome sequencing analysis revealed two novel deep-intronic variants in *PCDH15*

Proband 071203, clinically diagnosed with USH1, remained genetically unresolved after initial testing through molecular inversion probe (MIPs) analysis for 108 IRD-associated genes [30]. The MIPs-panel did not include the *PCDH15* gene, so we selected this case for WGS. Subsequent WGS analysis revealed a likely pathogenic start-loss variant in *PCDH15* (NM_033056.4, MANE Plus Clinical), c.1A>G, p.(Met1?) and a novel deep-intronic variant c.3983+3635A>G (p.(?)) in intron 29. For this variant we predicted the potential inclusion of a 61 bp pseudoexon (NM_033056.4, MANE Plus Clinical) based on the SpliceAI delta scores for acceptor gain (Δ0.63, position +65 bp from the variant-site) and donor gain (Δ0.69, position +5 bp from the variant-site) (**Figure 1A**). Both variants are absent from the control populations in gnomAD (Supplemental table 5). Sanger sequencing confirmed that the two *PCDH15* variants identified in this proband are *in trans*. The c.1A>G variant was found heterozygous in the father, the c.3983+3635A>G variant was found heterozygous in the mother, and neither variant was detected in the unaffected sibling.

**Figure 1.**
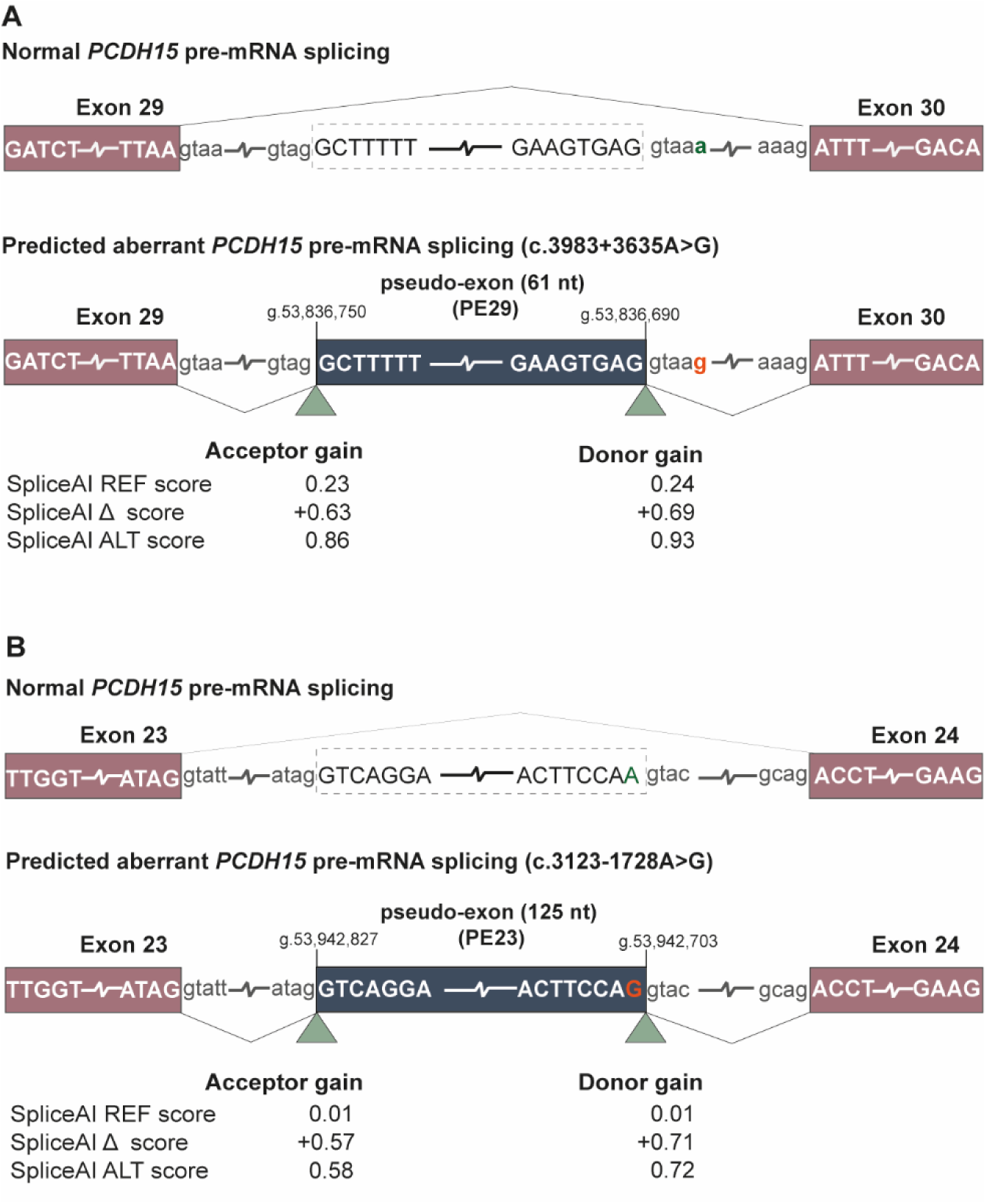
Schematic representation of normal and predicted aberrant pre-mRNA splicing in *PCDH15*. Normal and aberrant pre-mRNA splicing of two novel deep-intronic variants in *PCDH15* (NM_033056.4, MANE Plus Clinical). The top image depicts canonical pre-mRNA splicing connecting exons 29 and 30 **(A)** or connecting exons 23 and 24 **(B)** of *PCDH15*. Exons are shown as solid red boxes and intronic sequences in dashed boxes. Bottom images depict hypothesized aberrant gene splicing for variant c.3983+3635A>G **(A)** and variant c.3123-1728A>G **(B).** Expected pseudoexons are depicted in solid blue boxes. Green triangles depict the position of acceptor and donor gain as predicted by SpliceAI.

The second proband (DNA15-18518) and their sibling (DNA13-02611), clinically diagnosed with atypical USH, remained genetically unresolved after exome sequencing. While no conclusive pathogenic variants were found, the analysis did reveal two missense variants in *PCDH15* (NM_033056.4, MANE Plus Clinical) c.2885G>T (p.(Arg962Leu)) and c.2581G>A (p.(Val861Met)), but these were classified variants as benign and variant of unknown significance, respectively. Subsequent WGS in DNA15-18518 revealed an additional variant in *PCDH15*: a novel deep-intronic variant, c.3123-1728A>G. Targeted Sanger sequencing also confirmed the presence of this variant in DNA13-02611. This deep-intronic variant is located in intron 23 and has a reported global allele frequency of 0.0000855 in gnomAD. SpliceAI indicated delta scores of Δ0.57 (position +124 nt from the variant-site) for an acceptor gain and Δ0.71 (position +0 nt from the variant-site) for a donor gain (**Figure 1B**). Based on these observations, we hypothesized the inclusion of a pseudoexon of 125 nt. Although all three variants were identified in both affected siblings, the phase of these variants could not be determined as DNA from additional family members was not available. Consequently, it remains unclear which of these variants are *in trans* and which or if the combination of variants may explain the clinical phenotype.

To clarify the contribution of c.3983+3635A>G and c.3123-1728A>G to disease, we further extended our analysis of these variants to a cohort of nine genetically unresolved cases monoallelic for a pathogenic variant in *PCDH15*. We confirmed the presence of variant c.3123-1728A>G in two out of nine probands (DNA20-15335 and DNA21-01352). A pathogenic splice acceptor altering variant, c.3374-2A>G (NM_033056.4, MANE Plus Clinical), was previously identified using exome sequencing in proband DNA20-15335, whereas in proband DNA21-01352 a pathogenic 2 bp insertion variant, c.1779_1780insGA; p.(Arg594Aspfs*27) (NM_033056.4, MANE Plus Clinical), was detected (**Supplemental Table 5**). Through segregation analysis, we confirmed for both probands that the newly identified c.3123-1728A>G occurs *in trans* to their *PCDH15* pathogenic alleles identified by exome sequencing.

## Deep-intronic variants c.3123-1728A>G and c.3983+3635A>G in *PCDH15* lead to the insertion of pseudoexons

To evaluate the effect of deep-intronic variants on *PCDH15* pre-mRNA splicing, minigene splice assays were performed. For variant c.3983+3635A>G we detected a single fragment that corresponds to the size of the aberrant transcript including the predicted pseudoexon in intron 29 (PE29) (**Figure 2A**). By subsequent sequence validation using Sanger sequencing, we confirmed the use of the splice acceptor and donor sites as predicted by SpliceAI. The inclusion of the 61-nt pseudoexon in the transcript leads to a frameshift and premature stop codon in exon 30 of *PCDH15,* p.(Phe1329Leufs*28), thereby resulting in the premature termination of protein translation.

**Figure 2.**
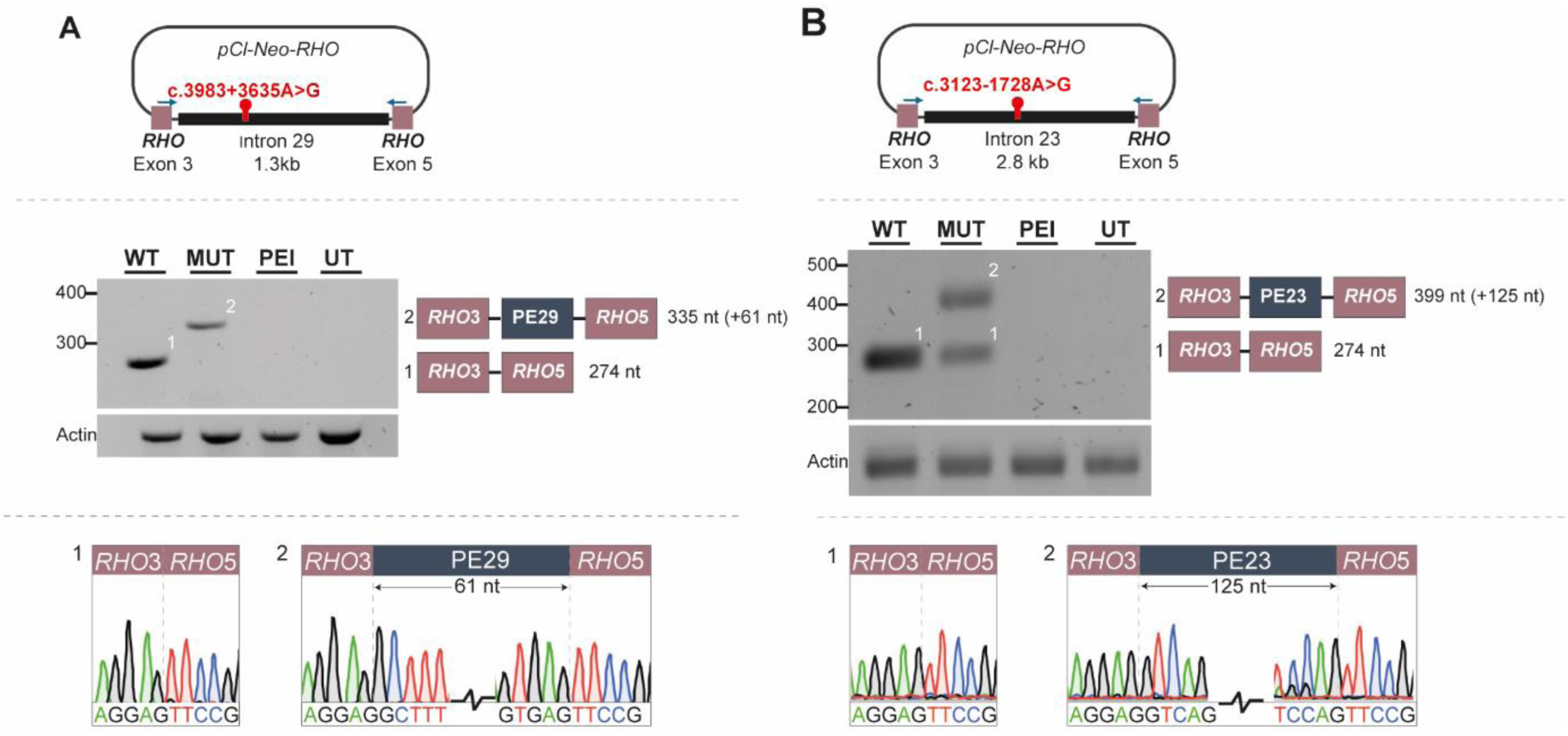
Minigene splice assay for c.3983+3635A>G (PE29) and c.3123-1728A>G (PE23) in HEK293T cells demonstrating a variant-induced splice defect in *PCDH15*. Top panels illustrate a schematic overview of the pCl-Neo-*RHO* minigene construct containing *rhodopsin* exons 3 and 5 and *PCDH15* intronic sequence harboring the deep-intronic variant, c.3983+3635A>G **(A)** and c.3123-1728A>G **(B).** Exons are depicted in red boxes, intronic sequences as black lines. **Mid panels** show RT-PCR transcript analyses derived from both wildtype (WT) and mutant (M) minigene constructs alongside negative controls PEI (PEI) and untransfected cells (UT). Distinct bands correspond to normal (1) or aberrantly (2) spliced transcripts. **Bottom panels** display Sanger sequencing chromatograms confirming aberrantly spliced minigene products.

For variant c.3123-1728A>G, we observed two fragments following gel electrophoresis, one fragment that corresponds to the size of the transcript including the predicted pseudoexon in intron 23 (PE23) and one fragment corresponding to the transcript without the inclusion of the pseudoexon (**Figure 2B**). Sanger sequencing confirmed that a 125-nt pseudoexon was observed in part of the transcripts, indicating a partial effect on pre-mRNA splicing for this variant in this minigene splice assay. Inclusion of the 125-nt pseudoexon in the transcript leads to a frameshift and premature stop codon in exon 24 of *PCDH15*, p.(Pro1042Serfs*46).

## Antisense oligonucleotides correct aberrant pre-mRNA splicing for two *PCDH15* **variants**

We designed an ASO-mediated splice correction strategy to rescue the aberrant inclusion of the pseudoexons resulting from the corresponding splice-altering variants. For each variant, two ASOs were designed to specifically block exonic splice enhancer motifs or splice donor/acceptor sites near or within the activated pseudoexon (**Figure 3A**).

**Figure 3.**
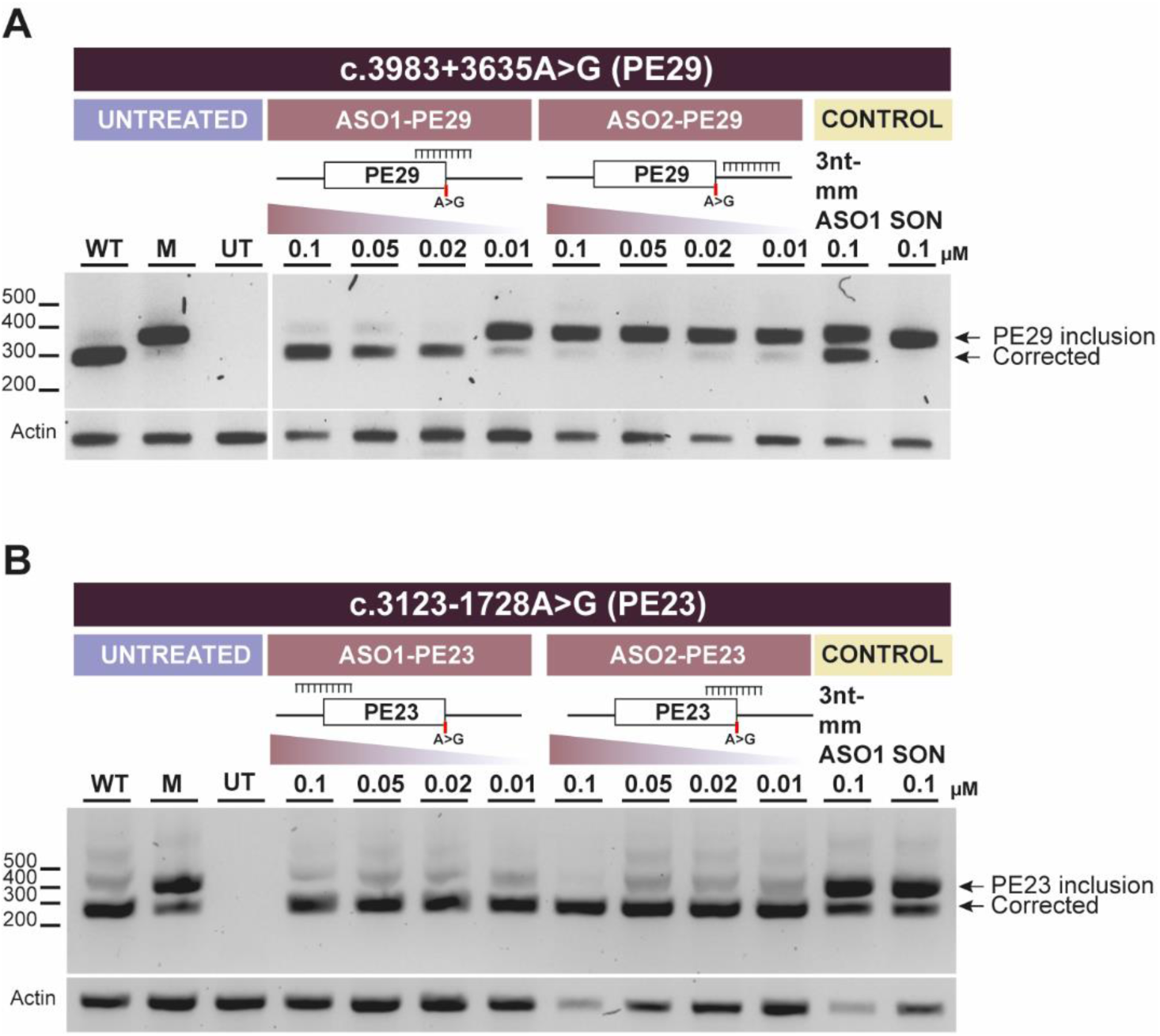
Antisense oligonucleotide (ASO) treatment for c.3983+3635A>G (PE29) and c.3123-1728A>G (PE23) in HEK293T cells using minigene constructs. RT-PCR and subsequent gel electrophoresis on mRNA of HEK293T cells co-transfected with ASO1 or ASO2 and either a wildtype or mutant minigene construct (A c.3983+3635A>G (PE29), B c.3132-1728A>G (PE23)). HEK293T cells were treated with ASO for 48 hours before harvesting. A three-nucleotide mismatch ASO (3nt-mm ASO) and a scrambled oligonucleotide (SON) were tested and were shown to have minimal effect on reducing the inclusion of the pseudoexons.

Through co-transfections of HEK293T cells with ASOs and matching minigene splice constructs, we determined that inclusion of PE29 resulting from c.3983+3635A>G was prevented by ASO1-PE29 with a minimally effective concentration of 0.02 µM, while ASO2-PE29 was unable to correct the splice defect at any of the concentrations used (**Figure 3A**). Transfection of the 3nt-mmASO of ASO1-PE29 resulted in a significantly reduced splice redirection as compared to the original ASO1-PE29, while transfection with the SON did not show any signs of splicing modulation, indicating that the splice correction observed for ASO1-PE29 is sequence-specific. For c.3123-1728A>G, both designed ASOs showed high splicing correction potential with all concentration used in the assay (0.01-0.1 µM) (**Figure 3B**).

## Antisense oligonucleotides correct aberrant *PCDH15* pre-mRNA splicing in patient-derived photoreceptor precursor cells

To further study the inclusion of PE23 in the *PCDH15* mRNA resulting from variant c.3123-1728A>G and to assess the splice correction potential of the designed ASOs in a patient-derived cellular model, we generated PPCs from induced pluripotent stem cells derived from proband DNA20-15335, who is compound heterozygous for the c.3123-1728A>G and the c.3374-2A>G variants in *PCDH15*. Next, we performed transcript analysis with primers in exon 23 and exon 25 flanking the deep-intronic variant (intron 23). The inclusion of PE23 was observed in this cellular model, confirming the results obtained from the *in vitro* minigene splice assay (**Supplemental Figure 1**).

In contrast to the ASO-mediated splice correction in HEK293T cells, treatment with ASO1-PE23 did not result in a clear splice correcting effect in PPCs, whereas ASO2-PE23 appeared as effective in correcting aberrant *PCDH15* splicing as in minigene splice assays (**Figure 4**). Through semi-quantitative analysis of fragment intensities, we determined that, upon treatment with ASO2-PE23, 95% of *PCDH15* transcripts lacked PE23 whereas in untreated or SON-treated PPCs only 60% of transcripts lacked PE23. Of note, both untreated and SON-treated PPCs showed an equal percentage of *PCDH15* transcripts lacking PE23 (∼60%), indicating that the effect of ASO2-PE23 in redirecting aberrant *PCDH15* splicing is indeed sequence-specific.

**Figure 4.**
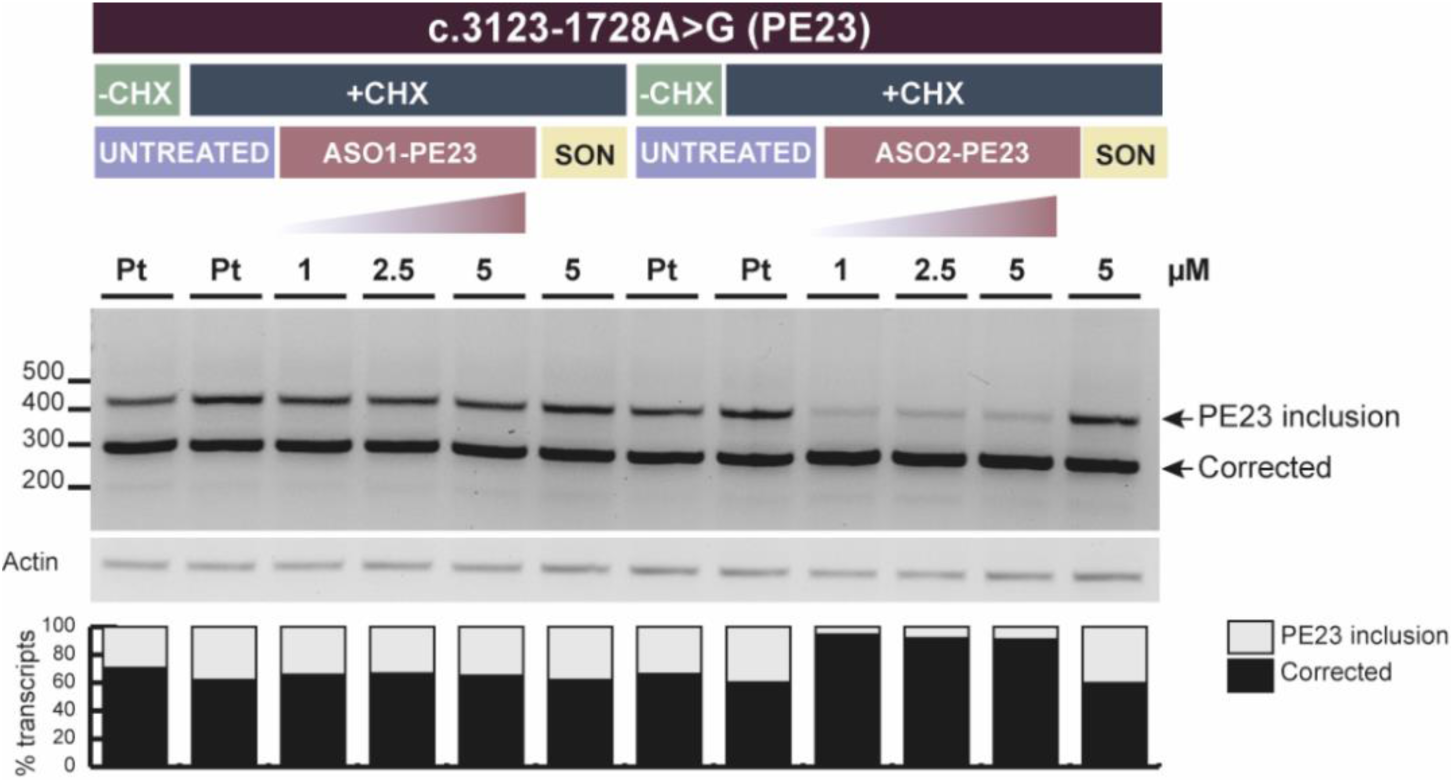
Antisense oligonucleotide (ASO) treatment of patient-derived photoreceptor precursor cells (PPC) harboring the heterozygous *PCDH15* c.3123-1728A>G (PE23) variant. RT-PCR and gel electrophoresis on patient (Pt) derived mRNA of PPCs harboring *PCDH15* variant c.3132-1728A>G (PE23) that were treated with two different ASOs at three different concentrations, treated with and without cycloheximide (CHX). Treatment with ASO2-PE23 resulted in a redirection of aberrant *PCDH15* splicing across all tested concentrations. Treatment with the scrambled oligonucleotide (SON) did not result in splicing redirection. Semi-quantitative analyses of fragment intensities are shown in a bar graph; black bars indicate the percentage of corrected transcript; grey bars indicate inclusion of PE23.

## Discussion

Bi-allelic pathogenic variants in *PCDH15* have been identified as the underlying cause of USH1, as well as for DFNB23. In this study, we identified two novel deep-intronic variants in *PCDH15*, c.3983+3635A>G and c.3123-1728A>G, that result in the inclusion of different pseudoexons. Following the identification of these novel splice-altering variants through WGS, we extended our analysis to a local cohort of nine probands with a clinical diagnosis of USH or non-syndromic deafness that were monoallelic for *PCDH15* variants. Via Sanger sequencing, we identified variant c.3123-1728A>G as a recurrent variant in two out of nine additional probands of this cohort. Variant c.3983+3635A>G was not identified in this cohort. The proband in which we identified variant c.3983+3635A>G is of Bukharan Jewish origin, and it is therefore not surprising that this variant was not found in a local cohort of Dutch patients.

To assess the effect of both deep-intronic variants on *PCDH15* pre-mRNA splicing, we employed an *in vitro* minigene splice assay using HEK293T cells. Results obtained from the *in vitro* minigene splice assay indicated that both variants indeed result in altered pre-mRNA splicing. While a minigene splice assay is usually a simple and cost-effective assay, it is primarily an effective first-line screening approach to functionally evaluate putative splice variants, but also has its limitations. First, the assay uses an artificial gene construct in which only a part of the gene of interest is cloned between two flanking *RHO* exons. In our case, the incorporation of flanking exonic *PCDH15* sequences was not feasible due to size constraints, as *PCDH15* introns 23 and 29 encompass 18,765 nt and 8,786 nt respectively. This could potentially affect the RNA folding of the *PCDH15* intronic sequences and subsequent binding of splicing factors. In addition, the assay makes use of a strong CMV promoter to overexpress a gene that is not endogenously expressed in the cell line and thus lacks the tissue specific splicing factors, such as those active in photoreceptor cells. These limitations may explain the observed discrepancy between HEK293T cells and the patient-derived PPCs in ASO experiments, where PE23-ASO2 was the only effective ASO in PPCs, while both ASOs seemed equally effective in HEK293T cells. This underscores the artificial nature of the minigene splice assay and the need to validate both splice defects and therapeutic interventions in cellular contexts that better reflect the patient’s genetic and gene regulatory landscape. This was also corroborated by the difference in the effect of *PCDH15* c.3132-1728A>G on pre-mRNA splicing seen in HEK293T cells (partial) and in patient-derived PPCs (almost complete), illustrating how the same variant behaves differently depending on the genetic and cellular context.

Previously, bi-allelic hypomorphic *PCDH15* alleles were linked to non-syndromic hearing loss whereas combinations with more severe and protein truncating variants were associated with USH1 [7]. This suggests that transcripts containing hypomorphic variants will result in the production of proteins with residual function sufficient for normal vision, but not for hearing. Variant c.3983+3635A>G was identified in 071203 with a clinical diagnosis of USH1. Variant c.3123-1728A>G was identified in mono-allelic probands DNA20-15335 and DNA21-01352 who both presented with severe bilateral hearing loss, but for whom we have no reliable data with respect to visual impairment and vestibular dysfunction of these probands, due to their young age. However, considering the effect of this variant on *PCDH15* pre-mRNA splicing, we consider that these variants explain the clinical phenotypes in these patients and recommend follow up of DNA20-15335 and DNA21-01352 to closely monitor development of visual complaints in the future.

Proband 071203 remained genetically unresolved after initial testing through MIPS analysis. At the time (2018), the MIPs panel covered 108 genes associated with non-syndromic IRD and did not include the *PCDH15* gene. For probands DNA20-15335 and DNA21-01352, monoallelic pathogenic *PCDH15* variants were identified through exome sequencing, but both probands remained genetically unresolved due to the lack of the identification of a (likely) pathogenic variant on the second allele. The *PCDH15* gene spans nearly 1 Mb with intron sizes up to 150 kb, providing ample space for a second pathogenic variant not being captured by exome sequencing. Moreover, deep-intronic pathogenic variants have been reported for multiple USH genes, including *PCDH15*. Indeed, through analysis of WGS data (071203) and via Sanger sequencing (DNA20-15335 and DNA21-01352) we now successfully completed the genetic diagnosis for these probands by the identification of deep-intronic variants. This highlights the diagnostic value of the genetic reanalysis of previously unresolved probands using technologies such as WGS, followed by a functional assessment of selected identified intronic variants of uncertain significance.

Identification of causative disease variants is crucial for affected individuals to become eligible for receiving future, often personalized, genetic therapies that are under development. Splice-altering variants provide promising targets for personalized RNA-based therapeutic strategies, such as ASO-mediated correction of splice defects. While the preclinical development and application of ASOs have been described for syndromic and non-syndromic IRDs such as Leber congenital amaurosis, Stargardt disease and Usher syndrome type 2a their use to therapeutically modulate *PCDH15* pre-mRNA splicing has not yet been described [31–35].

In this study, we therefore explored the use of ASOs to redirect aberrant *PCDH15* splicing caused by the identified deep-intronic variants and demonstrated that at least one of the ASOs designed per target effectively corrected aberrant pre-mRNA splicing events. Our data offer a foundation for further development towards a personalized therapeutic strategy. Future steps to enable an eventual clinical application should follow established guidelines, mainly related to an experimental N-of-1 treatment, and include additional functional studies on protein restoration and pathology in a suitable cell model, such as patient-derived organoids, and toxicity studies (both *in vitro* and *in vivo*) [36].

While ongoing studies have shown that by utilizing *mini-PCDH15* gene augmentation hearing loss can be rescued in a mouse model of USH1F when treated before postnatal day 5, which equals an *in utero* stage of development in man, it is unlikely that this approach could be translated into a treatment option to improve the hearing deficits of children with USH1, due to the lack of a therapeutic window of opportunity [37]. In contrast, the first signs of RP in these individuals typically present as night blindness during puberty followed by a slowly progressive peripheral vision loss. This provides a window of opportunity to intervene with personalized treatments, such as mutation-specific ASO-based splicing modulation, to rescue aberrant gene splicing and thereby halting the deterioration of vision. This underlines the importance of genetic testing and diagnosis to become eligible for receiving future genetic therapies.

In conclusion, this study identified two novel deep-intronic variants within the *PCDH15* gene which are both amenable to ASO-mediated splice correction. While ASOs have not yet been described as a therapeutic treatment option for *PCDH15*-associated disease, the findings in this study are promising and suggest the potential applicability of ASO-based therapies for a broader spectrum of patients with similar splice-altering, deep-intronic variants within the same gene.

## Acknowledgements

The authors want to thank all the patients and their family members for their participation in this study. The authors would like to thank the Department of Human Genetics and the Radboud Genome Technology Center for infrastructural and computational support. The authors wish to thank the Radboudumc Stem Cell Technology Center (https://www.radboudumc.nl/en/research/radboud-technology-centers/stem-cells) for reprogramming and characterizing the patient line. We would also like to thank Ellen Blokland, Saskia van der Velde-Visser and Marlie Jacobs-Camps for DNA sample preparation and administration.

## Data availability statement

Data are available upon reasonable request. All other genome sequencing data are subject to controlled access because they may compromise the privacy of research participants. These data may become available upon a data transfer agreement approved by the local ethics committee and can be obtained after contacting the corresponding author (S.R.) upon request.

## Supplementary figure

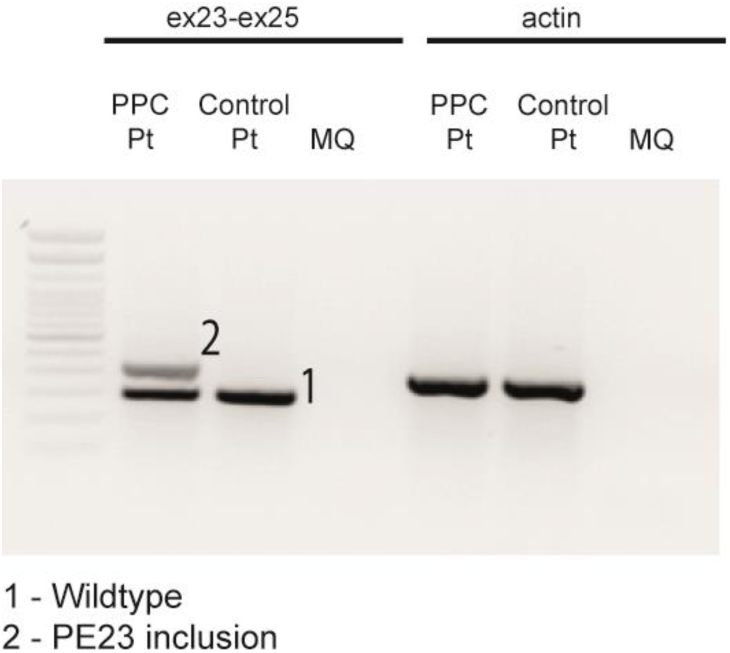
Figure S1.

## Supplemenal tables

**Supplemental table 1.**
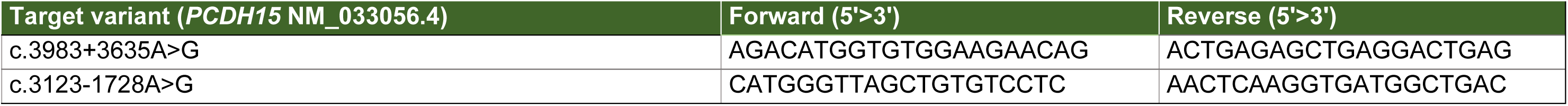
Sequences of primers used to assess the presence of novel *PCDH15* variants in an extended cohort.

**Supplemental table 2.**
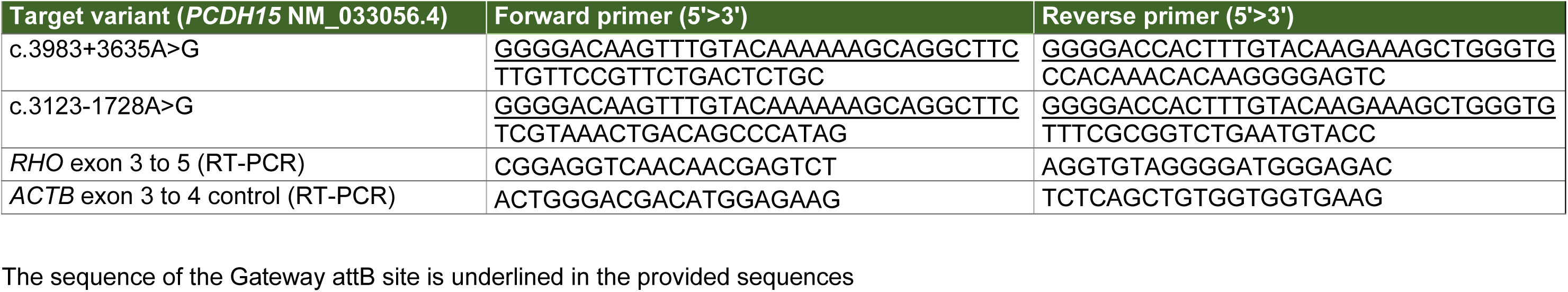
Sequences of primers used to generate minigene splice constructs.

**Supplemental table 3.**
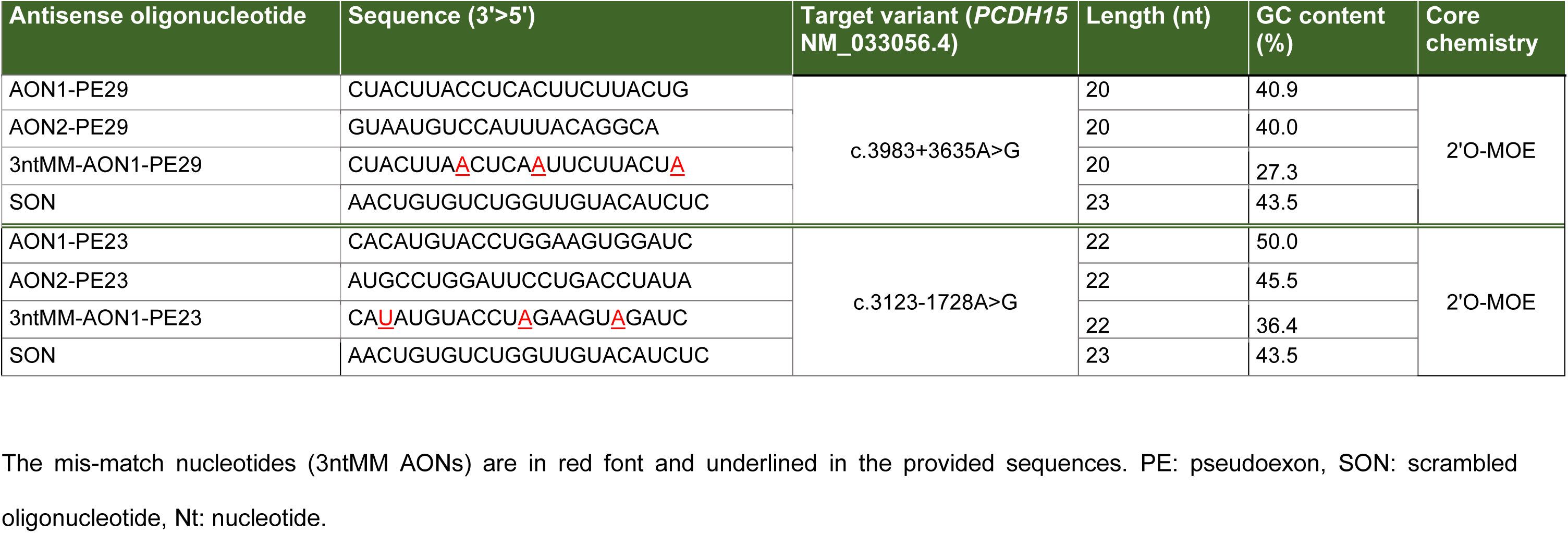
Sequences of antisense oligonucleotides used in this study.

**Supplemental table 4.**
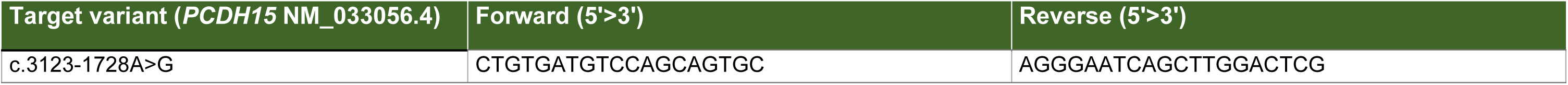
Sequences of primers used to assess the presence of the splice-altering variant in photoreceptor precursor cells.

**Supplemental table 5.**
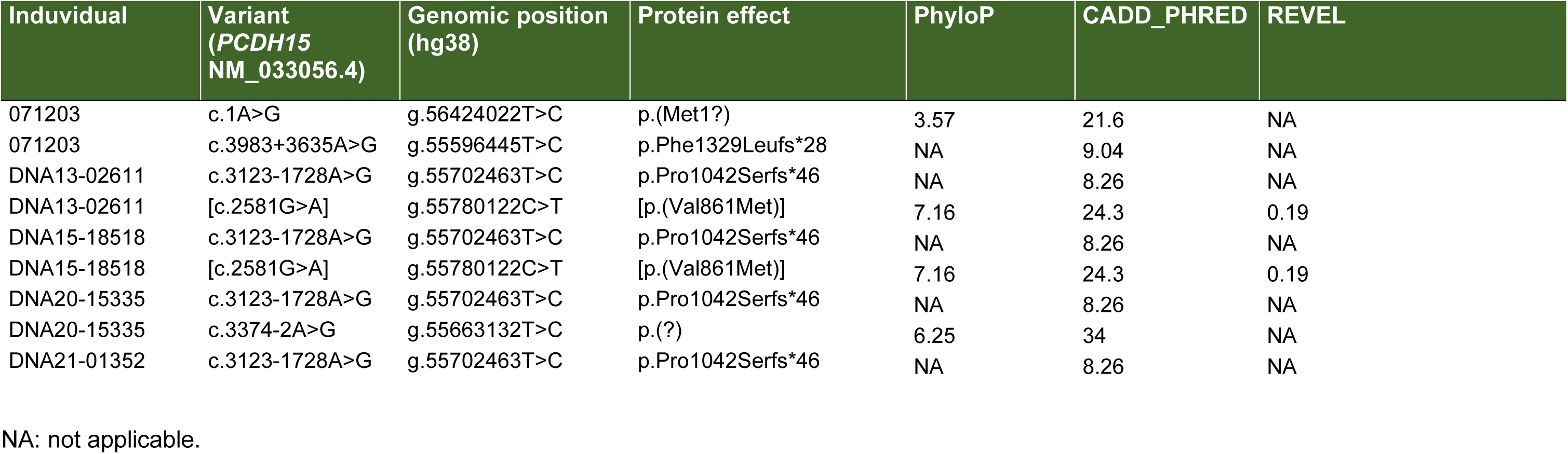
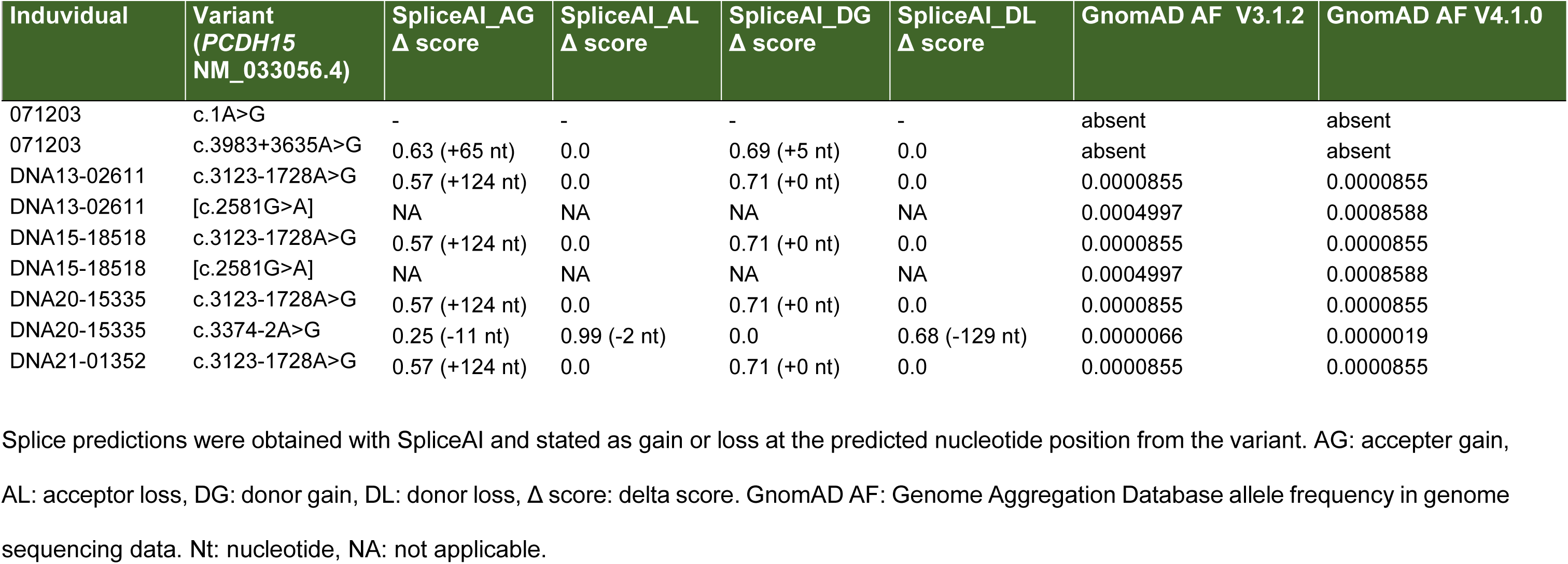
Overview of variants identified in this study.

