## Supplemental figure and tables for "Genome sequencing reveals novel pathogenic deep-intronic *PCDH15* variants, amenable to antisense oligonucleotide-based splice correction"

### Supplementary figure

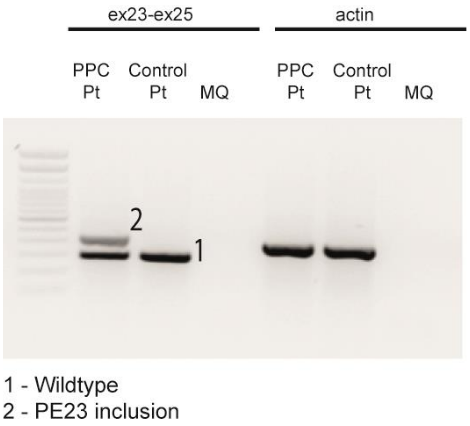


**Figure S1**

### Supplemenal tables

**Supplemental table 1.** Sequences of primers used to assess the presence of novel *PCDH15* variants in an extended cohort

| Target variant (*PCDH15* NM_033056.4) | Forward (5'>3') | Reverse (5'>3') |
|---|---|---|
| c.3983+3635A>G | AGACATGGTGTGGAAGAACAG | ACTGAGAGCTGAGGACTGAG |
| c.3123-1728A>G | CATGGGTTAGCTGTGTCCTC | AACTCAAGGTGATGGCTGAC |

**Supplemental table 2.** Sequences of primers used to generate minigene splice constructs.

| Target variant (*PCDH15* NM_033056.4) | Forward primer (5'>3') | Reverse primer (5'>3') |
|---|---|---|
| c.3983+3635A>G | GGGGACAAGTTTGTACAAAAAAGCAGGCTTC<br>TTGTTCCGTTCTGACTCTGC | GGGGACCACTTTGTACAAGAAAGCTGGGTG<br>CCACAAACACAAGGGGAGTC |
| c.3123-1728A>G | GGGGACAAGTTTGTACAAAAAAGCAGGCTTC<br>TCGTAAACTGACAGCCCATAG | GGGGACCACTTTGTACAAGAAAGCTGGGTG<br>TTTCGCGGTCTGAATGTACC |
| *RHO* exon 3 to 5 (RT-PCR) | CGGAGGTCAACAACGAGTCT | AGGTGTAGGGGATGGGAGAC |
| *ACTB* exon 3 to 4 control (RT-PCR) | ACTGGGACGACATGGAGAAG | TCTCAGCTGTGGTGGTGAAG |

The sequence of the Gateway attB site is underlined in the provided sequences

**Supplemental table 3.** Sequences of antisense oligonucleotides used in this study.

| Antisense oligonucleotide | Sequence (3'>5') | Target variant (*PCDH15* NM_033056.4) | Length (nt) | GC content (%) | Core chemistry |
|---|---|---|---|---|---|
| AON1-PE29 | CUACUUACCUCACUUCUUACUG | c.3983+3635A>G | 20 | 40.9 | 2'O-MOE |
| AON2-PE29 | GUAAUGUCCAUUUACAGGCA | | 20 | 40.0 | |
| 3ntMM-AON1-PE29 | CUACUUAACUCAAUUCUUACUA | | 20 | 27.3 | |
| SON | AACUGUGUCUGGUUGUACAUCUC | | 23 | 43.5 | |
| AON1-PE23 | CACAUGUACCUGGAAGUGGAUC | c.3123-1728A>G | 22 | 50.0 | 2'O-MOE |
| AON2-PE23 | AUGCCUGGAUUCCUGACCUAUA | | 22 | 45.5 | |
| 3ntMM-AON1-PE23 | CAUAUGUACCUAGAAGUAGAUC | | 22 | 36.4 | |
| SON | AACUGUGUCUGGUUGUACAUCUC | | 23 | 43.5 | |

The mis-match nucleotides (3ntMM AONs) are in red font and underlined in the provided sequences. PE: pseudoexon, SON: scrambled oligonucleotide, Nt: nucleotide.

**Supplemental table 4.** Sequences of primers used to assess the presence of the splice-altering variant in photoreceptor precursor cells.

| Target variant (*PCDH15* NM_033056.4) | Forward (5'>3') | Reverse (5'>3') |
|---|---|---|
| c.3123-1728A>G | CTGTGATGTCCAGCAGTGC | AGGGAATCAGCTTGGACTCG |

**Supplemental table 5**. Overview of variants identified in this study

| Induvidual | Variant (*PCDH15* NM_033056.4) | Genomic position (hg38) | Protein effect | PhyloP | CADD_PHRED | REVEL |
|---|---|---|---|---|---|---|
| 071203 | c.1A>G | g.56424022T>C | p.(Met1?) | 3.57 | 21.6 | NA |
| 071203 | c.3983+3635A>G | g.55596445T>C | p.Phe1329Leufs*28 | NA | 9.04 | NA |
| DNA13-02611 | c.3123-1728A>G | g.55702463T>C | p.Pro1042Serfs*46 | NA | 8.26 | NA |
| DNA13-02611 | [c.2581G>A] | g.55780122C>T | [p.(Val861Met)] | 7.16 | 24.3 | 0.19 |
| DNA15-18518 | c.3123-1728A>G | g.55702463T>C | p.Pro1042Serfs*46 | NA | 8.26 | NA |
| DNA15-18518 | [c.2581G>A] | g.55780122C>T | [p.(Val861Met)] | 7.16 | 24.3 | 0.19 |
| DNA20-15335 | c.3123-1728A>G | g.55702463T>C | p.Pro1042Serfs*46 | NA | 8.26 | NA |
| DNA20-15335 | c.3374-2A>G | g.55663132T>C | p.(?) | 6.25 | 34 | NA |
| DNA21-01352 | c.3123-1728A>G | g.55702463T>C | p.Pro1042Serfs*46 | NA | 8.26 | NA |

NA: not applicable.

**Supplemental table 5**. Overview of variants identified in this study (continued)

| Induvidual | Variant (*PCDH15* NM_033056.4) | SpliceAI_AG Δ score | SpliceAI_AL Δ score | SpliceAI_DG Δ score | SpliceAI_DL Δ score | GnomAD AF V3.1.2 | GnomAD AF V4.1.0 |
|---|---|---|---|---|---|---|---|
| 071203 | c.1A>G | - | - | - | - | absent | absent |
| 071203 | c.3983+3635A>G | 0.63 (+65 nt) | 0.0 | 0.69 (+5 nt) | 0.0 | absent | absent |
| DNA13-02611 | c.3123-1728A>G | 0.57 (+124 nt) | 0.0 | 0.71 (+0 nt) | 0.0 | 0.0000855 | 0.0000855 |
| DNA13-02611 | [c.2581G>A] | NA | NA | NA | NA | 0.0004997 | 0.0008588 |
| DNA15-18518 | c.3123-1728A>G | 0.57 (+124 nt) | 0.0 | 0.71 (+0 nt) | 0.0 | 0.0000855 | 0.0000855 |
| DNA15-18518 | [c.2581G>A] | NA | NA | NA | NA | 0.0004997 | 0.0008588 |
| DNA20-15335 | c.3123-1728A>G | 0.57 (+124 nt) | 0.0 | 0.71 (+0 nt) | 0.0 | 0.0000855 | 0.0000855 |
| DNA20-15335 | c.3374-2A>G | 0.25 (-11 nt) | 0.99 (-2 nt) | 0.0 | 0.68 (-129 nt) | 0.0000066 | 0.0000019 |
| DNA21-01352 | c.3123-1728A>G | 0.57 (+124 nt) | 0.0 | 0.71 (+0 nt) | 0.0 | 0.0000855 | 0.0000855 |

Splice predictions were obtained with SpliceAI and stated as gain or loss at the predicted nucleotide position from the variant. AG: accepter gain, AL: acceptor loss, DG: donor gain, DL: donor loss, Δ score: delta score. GnomAD AF: Genome Aggregation Database allele frequency in genome sequencing data. Nt: nucleotide, NA: not applicable.
